# Progressing adaptation of SARS-CoV-2 to humans

**DOI:** 10.1101/2020.12.18.413344

**Authors:** Tomokazu Konishi

## Abstract

The second and subsequent waves of coronavirus disease 2019 (COVID-19) have caused problems worldwide ^1^. Here, using an objective analytical method ^2^, we present the changes that occurred in the severe acute respiratory syndrome coronavirus 2 (SARS-CoV-2), the causative virus of COVID-19, over time. The virus has mutated in three major directions, resulting in three groups to date. Analysis of the basic structure of the group of viruses was completed by April and shared across all continents. However, the virus continued to mutate independently in each country after the borders were closed. In particular, the virus mutated before the occurrence of the second and subsequent peaks. It seems that the mutations conferred higher infectivity to the virus, because of which the virus overcame previously effective protection and caused second waves of the disease. Currently, each country may possess such a unique, stronger variant. Some of them slowly entered other countries and caused epidemics. These viruses could also serve as sources of further mutations by exchanging parts of the genome, which could create variants with superior infectivity.

## Introduction

The evolutionary trajectory of an evolved virus is difficult to analyse from nucleotide sequences because the data have complex multivariate structures with numerous dimensions ^2^. Phylogenetic trees have long been used to present relationships in among sequences ^3^ and many studies on severe acute respiratory syndrome coronavirus 2 (SARS-CoV-2) ^4^ have employed them ^5–8^, but they have two drawbacks. One is the decisive lack of falsifiability that ruins objectivity ^9^, and the other is a lack of generality that makes it difficult to integrate with other data sources.

Here, an analysis was performed using a more objective method, i.e., principal component analysis (PCA) ^10^. PCA runs a singular value decomposition on a sequence described as a Boolean vector and obtains the principal component (PC) for the sample and base in parallel ^11^. PCA finds common directions among a dataset and represents them as independent axes. These axes have an order; PC1 is common to most data, and the lower the contribution to differences in the data, the lower the order. The sample or base is presented on the axis according to its characteristics. As shown in the current study, SARS-CoV-2 data were represented by two axes as a whole. The lower axes represented mutations found in smaller areas. The components were scaled and compared with those of the influenza virus ^12^ as needed. In addition, the results were integrated with the date and location of collection.

## Results and Discussion

### Overview of PCA

PCs 1 and 2 formed a triangle comprising four groups that were temporarily termed as 0-3 (Fig. 1A). All groups were found on all continents. In contrast to PC1 and PC2 showing the overall situation, the PC3 axis and the lower-order axes showed differences that relate to only a small number of the samples (Fig. 1B and Extended Data Fig. 1), with smaller contributions to differences in the data (Extended Data Fig. 2); many of them were detected in limited countries after April (comparing Extended Data Fig. 1 and 3). Such mutations occurred since April when countries began to restrict the movement of people across national borders. The overview of Fig. 1A was completed at the end of March (Fig. 1C). Although group 3 once appeared worldwide, it has been contained to date (Fig. 1D).

**Fig. 1:**
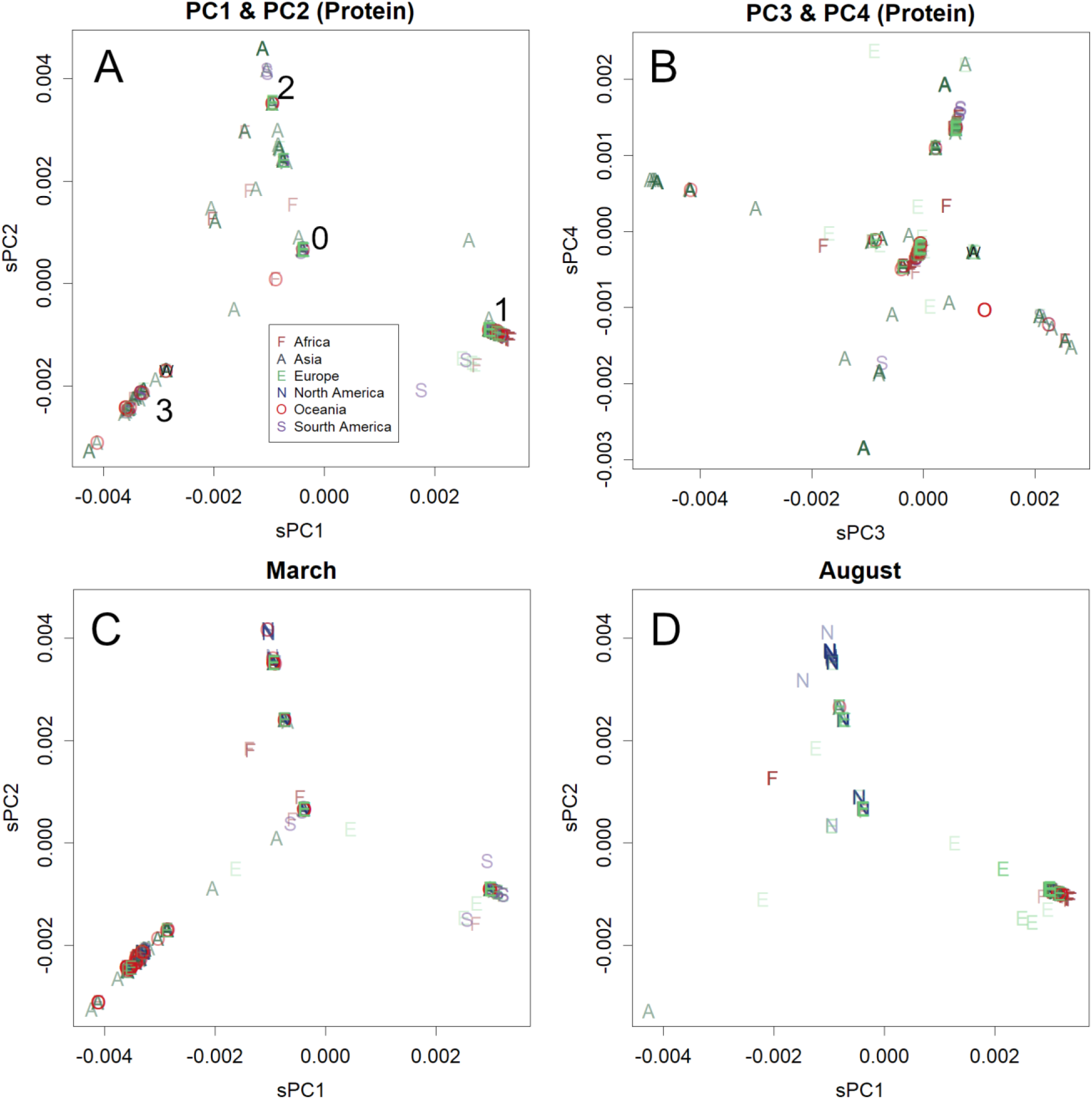
Principal components of SARS-CoV-2 amino acid sequences reported until December 1, 2020. One hundred samples are randomly selected from a continent and presented. **A**. PC1 and PC2. Numbers are given for a temporal explanation of the groups. **B**. PC3 and PC4. The earliest variants are indicated by the letter W. The numbers of samples that are separated by the axes are reduced in the lower-order axes (Extended Data Fig. 2 and 3). **C** and **D**. PC1 and PC2 at the end of March and from August 2020. A maximum of 100 samples is selected from a continent and presented. The components are scaled by the length of sequences; squares of the values are correlated with the expectation of differences per base (Supplementary Information).

These features were similar for both RNA and protein (Extended Data Fig. 1 and 4). The rate of missense mutations, which affects amino acid sequence, was 63% (Extended Data Fig. 5). This is a fairly high rate; rather, null mutations are expected to be 61% ^13^ even though some differences caused by ignoring damaging mutations have been identified here. In fact, in the case of H1N1 influenza virus from 2001 to 2003 in the United States, the rate was 22%. This suggested that SARS-CoV-2 was under selective pressure to alter its protein structure.

The viral groups causing the epidemics changed over time (Fig. 2A). The second and third waves in each country were caused by new variants of the virus, which showed different patterns at lower PCs (Fig. 2). In England, the second wave was caused by a variant of the Group 0, which has since spread to many countries in Europe (Extended Data Fig. 6, panels A-I). The third wave occurred due to another variant of Group 1, H69-V70 (Fig. 2C) ^14,15^.

**Fig. 2:**
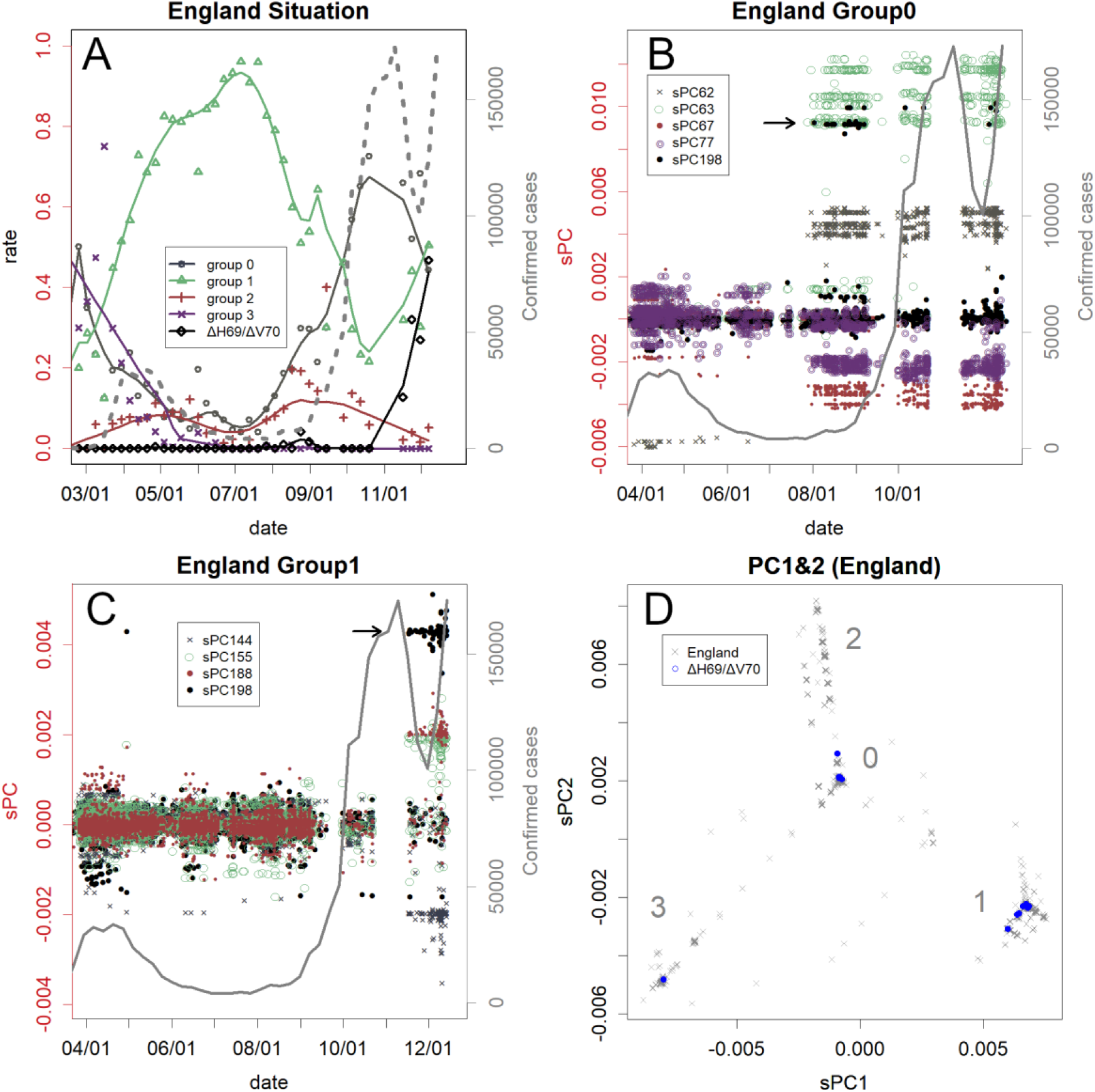
Mutations found on PCA and the number of patients in England. **A**. Changes in rates in each group. The dotted grey line shows the confirmed cases. The second and third winds were caused mainly by group 0 and 1 variant, respectively. **B**. PC fingerprints of group 0. Prior to the second wind, the pan-European variant appeared. PC198 (black dots) higher than the black arrow indicates the H69-V70 group 0 variant. **C**. PC fingerprints of group 1. Prior to the third wind, H69-V70 group 1 variant (black arrow) appeared. D. H69-V70 variants found in England (blue).

The H69-V70 variant in the UK ^14,15^ actually involves multiple mutations (Extended Data Table 2). The deletion itself has occurred spontaneously in multiple groups (Fig. 2D); the epidemic in the UK is caused by a group 1 variant, while those in Germany and Denmark are caused by group 0. There were other group 1 variants without the deletion in the UK, which would be the ancestral variants. The group 1 H69-V70 variant in the UK is growing at the same or faster rate than the pan-European variant by replacing it (Fig. 2A). These results show that this variant could be more infectious than the pan-European variant.

A group 2 variant is quickly spreading in South Africa (Extended Data Fig. 6Q and 6R). As it had not appeared until October, the axes of PCA had not trained for the variant; hence, the figure was made by renewing the axes. This is a novel variant that contains many mutations concentrated in the spike protein, and the magnitude of mutation is deemed comparable to that of influenza H1N1 hemagglutinin between sequential peaks (Table 1). Group 2 variants have been negligible in South Africa (Extended Data Fig. 6Q), and no variant was considered ancestral. Perhaps the epidemic variant was from areas where samples are rarely sequenced. Any region in the world can produce a new highly infectious variant, with potential to cross national borders. Therefore, measures such as vaccines are urgently needed on a global scale.

**Table 1.** The rate of mutated amino acid residues during the indicated years and location. The numbers show mutations per 1000 amino acid residues. Results of the H1N1 influenza virus hemagglutinin protein observed in the peak seasons, and the whole genome of coronavirus reported in the peaks are shown. (2008–2009) indicates the difference between strains before 2009 and the pdm09 influenza virus. Max (PC1, PC2) is the difference between the most diverged SARS-CoV-2 groups in Fig. 1A. The England (group 1) and South Africa (group 2) are the new variants presently spreading rapidly. The England variant (Group 1) and the South African one (Group 2) are the new viral variants spreading at a rapid rate currently.

|  |  |
| --- | --- |
| Influenza H1N1 | /1000 |
| USA, 2001-2003 | 32 |
| New Zealand, 2001-2005 | 15 |
| (2008-2009) | 203 |
| USA, 2009-2014 | 15 |
| USA, 2014-2016 | 7.1 |
| Thailand, 2009 - 2010/2/1 | 0 |
| Thailand, 2010/2/1 - 10/10 | 2.5 |
| SARS-CoV-2 | /1000 |
| Austraria (group 1), 4/1 - 8/20 | 0.5 |
| France (group 2), 6/5 - 9/20 | 1 |
| England (group 0), 3/30-8/15 | 1.1 |
| South Africa (group 1), 5/30-8/3 | 0.8 |
| USA (group 2), 3/30-8/15 | 0.4 |
| England (group 1), Entire | 1.5 |
| South Africa (group 2), Spike | 10.6 |
| South Africa (group 2), Entire | 2.6 |

In many countries, mutations were detected several weeks prior to the peaks (Fig. 2 and Extended Data Fig. 6). This suggested that the mutations were the cause of the peak. In any case, the variants prevalent at those peaks should certainly be more infectious than they were before the mutation, so the variants were replaced. The higher infectivity may have weakened the previously effective protections from the virus. Altered residues between the peaks are shown in Extended Data Table 1; because PCA showed differences among samples and bases in parallel, PCs for bases may help identify those differences. It may be possible to predict the next wave by monitoring the mutations. However, only a limited number of countries continue sequencing the virus; many have reported only the first peak. However, the second and subsequent peaks are caused by different variants (Fig. 2 and Extended Data Fig. 6).

### Magnitude of mutations

Influenza H1N1 is highly infectious in humans ^16^. It mutates continuously but does not appear in the same area during two sequential years. It is prevalent every few years because it takes time to mutate enough to survive against herd immunity in the area. However, the magnitude of changes in the meantime varies (Table 1). Perhaps, the part of the sequence that needs to be modified to escape herd immunity varies from case to case. However, the changes in coronavirus before and after the peaks were much smaller (Table 1). The differences seem to be smaller than that needed to obtain another peak of seasonal influenza, although this does not guarantee safety from exceptional reinfections ^17,18^. A possible exception is the one that presently causes an epidemic in South Africa.

A case of pdm09 influenza virus that likely corresponded to SARS-CoV-2 mutations was observed in Thailand from 2009 to 2010 (Extended Data Fig. 7) ^19^. The 2009 pandemic was subdued in this tropical country, and three peaks were subsequently identified toward 2010. At the second peak, only the nucleotide sequence changed, and at the third peak, the hemagglutinin protein changed (Table 1) with a high rate of missense mutations (54%). However, these changes were not highly associated with herd immunity because most of the population was not yet infected. These changes might be caused by the adaptation of pdm09—originally a swine strain—to humans.

The influenza virus genome continues to mutate and not only in the part that is directly associated with infection ^12^. This property may be the reason it has repeatedly deceived herd immunity for decades, allowing influenza to remain as a pandemic.

In contrast, mutations in SARS-CoV-2 were not uniform; rather, some smaller open reading frames (ORFs) such as E, M, and ORF6 were preserved (Fig. 3). These could have adapted to humans, but they are also conserved in various coronaviruses ^20^. If they remain conserved, they could be subjected to herd immunity, extending the period between future epidemics. Evidently, this does not mean that these ORFs do not change forever; the rate of change is just slower.

**Fig. 3.**
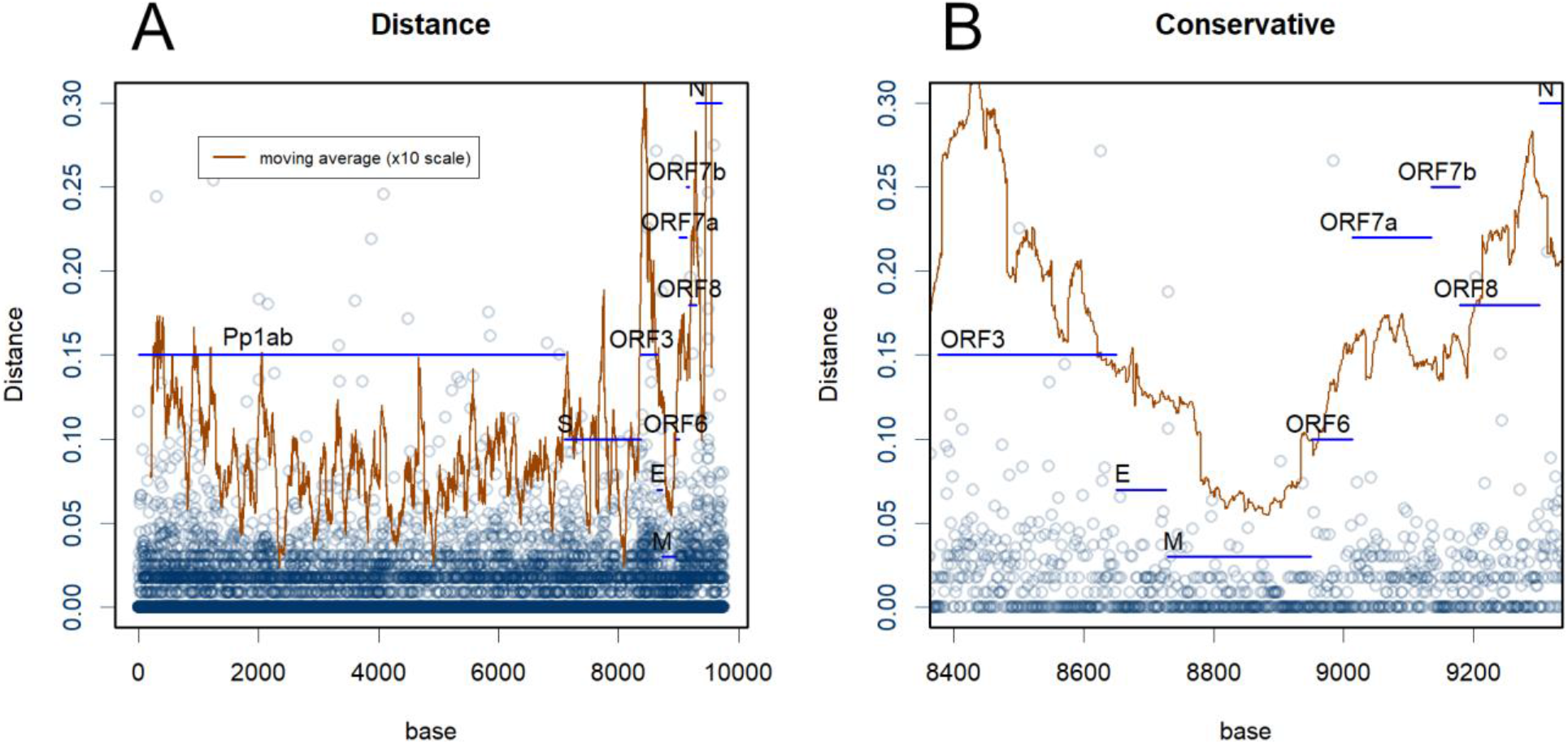
**A:** Mean distances to the sample mean (protein). The square of the distance reflects the expected mutation occurrence at the residue. The brown line indicates the moving average of 100 residues (×10 scale). **B.** Magnified view of the conserved region.

The strain responsible for the pandemic has mutated rapidly. As of March, the bases of PCs 1 and 2 were formed, and each continent harboured the same set of mutated variants (Fig. 1). Since then, human movement has been restricted, and each country accumulated its own variants. In this process, the force that caused the mutations was probably adaptation to humans (Table 1). It caused changes in both codon usage and protein structure (Extended Data Fig. 1 and 4), but the rate of missense mutations was high (Extended Data Table 1).

Some particularly infectious variants of the virus crossed borders and became the cause of the second wave in many countries (Extended Data Fig. 6). A Group 0 variant, which has caused the epidemic in Spain in July, was detected in other EU countries about a month later, leaving the same fingerprints in the PCs (panels B, D, F, I, and L). In countries other than France, this strain formed second waves (Fig. 2 and Extended Data Fig. 6). In France, a Group 2 variant was prevalent at that time (Extended Data Fig. 6H); currently, this pan-European variant is spreading at a rapid rate (panel I). The pan-European variant has also reached the USA, Japan, and Australia; however, the sequences of the epidemic variant are unknown in Japan and in the USA, though these countries are amidst waves of infection. Australia seems to have succeeded in controlling this variant (panel O).

The unique variants in each country would serve as the source of newer variants, which are more infectious than the previous variants; therefore, they replaced the mode (Fig. 2 and Extended Data Fig. 6). If countries had not closed their national borders, many of their variants could have been replaced by other more powerful variants, as in the case of group 3. However, the unique variants survived in a narrower area; the variant that is causing the epidemic in South Africa would be an example. International events, such as the Olympic Games, which attract people in large numbers, will serve as an exchange market for these viruses. To make matters worse, coronavirus could acquire shift-type mutations, resulting in a mixture of genomes ^20^. When an exported strain is mixed with a domestic strain, a new strain is created. If each of the unique mutations works independently (otherwise, replacement of the older variants with these variants would not be beneficial), the accumulation of the mutations can result in the production of more powerful viruses with cherry-picked the surviving unique mutations.

The epidemics showed much variation among the countries (Extended Data Fig. 6). A Group 2 variant that caused the second wave in the USA (panels M and N) migrated to Australia a few months later (panel P). However, in contrast to the USA where the epidemic did not settle down, Australia quickly suppressed the second wave. Furthermore, New Zealand did not even let an epidemic occur. It should be noted that Sweden, which did not enforce a lockdown, could not control the pan-European variant (panels K and L). Before this variant, the Swedish measures had worked well and the infection was avoided simply by social distancing. However, the pan-European variant was so infectious that it negated those defenses. The responsive measures of various countries should be compared in detail for future pandemic.

## Notes

https://figshare.com/articles/dataset/dx_doi_org_10_6084_m9_figshare_6025748/6025748

### Competing Interest Statement

The authors have declared no competing interest.

### Summary of Updates

The second and subsequent waves of coronavirus disease 2019 (COVID-19) have caused problems worldwide. Here, using an objective analytical method, we present the changes that occurred in the severe acute respiratory syndrome coronavirus 2 (SARS-CoV-2), the causative virus of COVID-19, over time. The virus has mutated in three major directions, resulting in three groups to date. Analysis of the basic structure of the group of viruses was completed by April and shared across all continents. However, the virus continued to mutate independently in each country after the borders were closed. In particular, the virus mutated before the occurrence of the second and subsequent peaks. It seems that the mutations conferred higher infectivity to the virus, because of which the virus overcame previously effective protection and caused second waves of the disease. Currently, each country may possess such a unique, stronger variant. Some of them slowly entered other countries and caused epidemics. These viruses could also serve as sources of further mutations by exchanging parts of the genome, which could create variants with superior infectivity.

## References

1 WHO. Coronavirus disease (COVID-19) Pandemic, <https://www.who.int/emergencies/diseases/novel-coronavirus-2019> (2020).

2 Konishi, T. et al. Principal Component Analysis applied directly to Sequence Matrix. Scientific Reports 9, 19297, doi:10.1038/s41598-019-55253-0 (2019).

3 Yang, Z. & Rannala, B. Molecular phylogenetics: principles and practice. Nat Rev Genet 13, 303–314, doi:10.1038/nrg3186 (2012).

4 WHO. WHO Coronavirus Disease (COVID-19) Dashboard, <https://covid19.who.int/> (2020).

5 Alm, E. et al. Geographical and temporal distribution of SARS-CoV-2 clades in the WHO European Region, January to June 2020. Euro Surveill 25, 2001410, doi:10.2807/1560-7917.ES.2020.25.32.2001410 (2020).

6 Zhou, P. et al. A pneumonia outbreak associated with a new coronavirus of probable bat origin. Nature 579, 270–273, doi:10.1038/s41586-020-2012-7 (2020).

7 Lam, T. T.-Y. et al. Identifying SARS-CoV-2 related coronaviruses in Malayan pangolins. Nature 583, 282–285, doi:10.1038/s41586-020-2169-0 (2020).

8 Murakami, S. et al. Detection and Characterization of Bat Sarbecovirus Phylogenetically Related to SARS-CoV-2. Emerging Infectious Diseases 26, 3025–3029 (2020).

9 Ellis, G. & Silk, J. Scientific method: Defend the integrity of physics. Nature 516, 321–323, doi:10.1038/516321a (2014).

10 Jolliffe, I. T. Principal Component Analysis. (Springer-Verlag 2002).

11 Konishi, T. Principal component analysis for designed experiments. BMC Bioinformatics 16 Suppl 18, S7, doi:10.1186/1471-2105-16-S18-S7 (2015).

12 Konishi, T. Re-evaluation of the evolution of influenza H1 viruses using direct PCA. Scientific Reports 9, 19287, doi:10.1038/s41598-019-55254-z (2019).

13 Wang, T. et al. Probability of phenotypically detectable protein damage by ENU-induced mutations in the Mutagenetix database. Nature Communications 9, 441, doi:10.1038/s41467-017-02806-4 (2018).

14 Bal, A. et al. Two-step strategy for the identification of SARS-CoV-2 variants co-occurring with spike deletion H69-V70, Lyon, France, August to December 2020. medRxiv, 2020.2011.2010.20228528, doi:10.1101/2020.11.10.20228528 (2020).

15 Volz, E. et al. Transmission of SARS-CoV-2 Lineage B.1.1.7 in England: Insights from linking epidemiological and genetic data. virological.org < https://virological.org/t/transmission-of-sars-cov-2-lineage-b-1-1-7-in-england-insights-from-linking-epidemiological-and-genetic-data/576>(2021).

16 CDC. Similarities and Differences *between Flu and COVID-19*, <https://www.cdc.gov/flu/symptoms/flu-vs-covid19.htm> (2020).

17 Gupta, V. et al. Asymptomatic Reinfection in 2 Healthcare Workers From India With Genetically Distinct Severe Acute Respiratory Syndrome Coronavirus 2. Clinical Infectious Diseases, doi:10.1093/cid/ciaa1451 (2020).

18 Edridge, A. W. D. et al. Seasonal coronavirus protective immunity is short-lasting. Nature Medicine 26, 1691–1693, doi:10.1038/s41591-020-1083-1 (2020).

19 WHO. Influenza surveillance outputs, <https://www.who.int/influenza/resources/charts/en/> (2020).

20 Konishi, T. Principal component analysis of coronaviruses reveals their diversity and seasonal and pandemic potential. PLoS ONE 15, e0242954, doi:10.1371/journal.pone.0242954 (2020).

21 Elbe, S. & Buckland-Merrett, G. Data, disease and diplomacy: GISAID’s innovative contribution to global health. Global challenges (Hoboken, NJ) 1, 33–46, doi:10.1002/gch2.1018 (2017).

22 Wright, E. S. DECIPHER: harnessing local sequence context to improve protein multiple sequence alignment. BMC Bioinformatics 16, 322, doi:10.1186/s12859-015-0749-z (2015).

23 R Core Team. R: A language and environment for statistical computing. (R Foundation for Statistical Computing, 2020).

