## Supplement for "Progressing adaptation of SARS-CoV-2 to humans"

**Supplementary Information for Changes in SARS-CoV-2 before the peak in each country, producing unique variants**

**Relationship between sPC and changed number of bases**

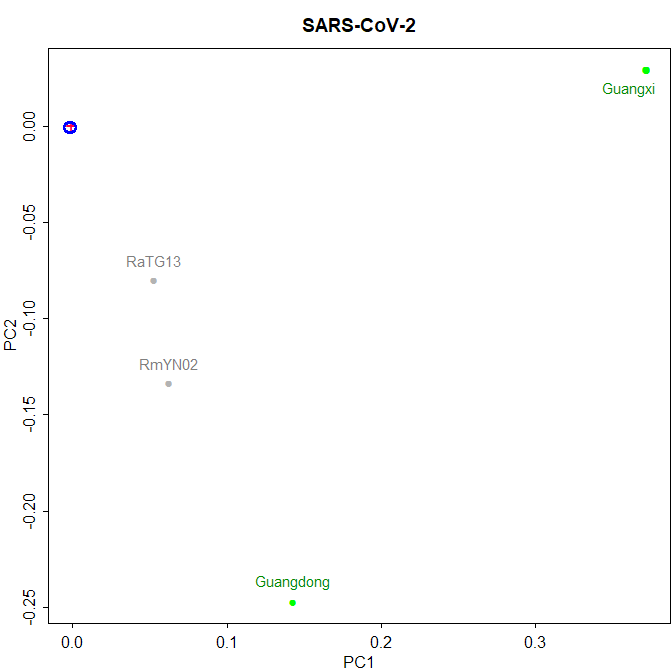
The principal component of a sample represents a part of the Euclidean distance from the mean sample, the square root of the sum of squares of the difference, $l=\sqrt{{\sum(\mathrm{sample}-\mathrm{mean})^{2}}/2}$ , where “sample” is the Boolean vector of a sample and “mean” is the arithmetic mean ^1^. The components were further scaled by nucleotide length owing to the generality of the values. Otherwise, because influenza virus and coronavirus are very different in terms of genome length, the results cannot be compared.

The correlation between base changes and the PCs is as follows. For example, in this figure, the Euclidean distance between humans (blue) and bat RaTG13 (grey) is 0.1. The figure shows the square root of the expected difference in each base (because the values are scaled). Therefore, the total difference was 0.1 ^ 2 × length = 280. This was only part of the total difference of 1100 because PCA identified many other axes and the distance was partitioned for other axes as well. If all differences that appear on all axes are summed, the total is 1100. The length of 280 is the difference that appeared in the two directions shown in the PC1 and PC2 axes.
