## Extended Data for "Progressing adaptation of SARS-CoV-2 to humans"

**Materials and Methods**

*PCA.* Nucleotide sequences were obtained from GISAID ^21^ on October 1, 2020. Only the complete sequences that did not contain N were selected. Sequences were aligned using DECHIPER ^22^. The sequences were converted to a Boolean vector and subjected to PCA ^2^. Sample PCs and sequence PCs are scaled based on the length of the sequence and the number of samples, respectively ^11^. All calculations were performed using R ^23^. The ID, acknowledgements, and scaled PCs of samples and bases can be downloaded from Figshare (dx.doi.org/10.6084/m9.figshare.6025748). The newest version of R code and the PCA axes are publicly available (https://github.com/TomokazuKonishi/Fast-calc-for-SARS-CoV-2). Using these methods, any sequence data can be represented on the same axes presented here, and the axes can also be updated with newer sequences. The relationship between PC values and the number of mutated bases is briefly explained in the Supplementary Information. Number of confirmed cases were obtained from WHO ^4^.

The axes were identified and used as follows. Because PCA axes are sensitive to sample bias, they were determined using 6,092 samples randomly selected from each continent in proportion to their population; 989 from Africa, 3636 from Asia, 610 from Europe, 476 from North America, 347 from South America, and 34 from Oceania; the number of sequences used from Africa was the maximum available; hence, the sampling was not random. For clarity, Fig. 1 presents only 100 samples from each continent, whereas Extended Data figures present approximately 1000 samples. The contributions of the axes are presented in Extended Data Fig. 2.

*Distances between samples from different peaks.* In influenza, only one strain has emerged over the years, with repeated mutations ^12^. Differences between peaks within the same strain were found as differences between the mean sequences from both peaks. In contrast, in coronavirus, the peak comprises several different variants, especially the first peak. Averaging will cancel the characteristics of these variants. To avoid this problem, the differences were estimated among those that have similar PC1 and PC2 and are in the same group as shown in Table 1, and the differences only appeared in the lower PCs.

*The rate of missense mutations.* In influenza, differences in both nucleotide and amino acid sequences were presumed as differences between the average peak sequences. For coronavirus, the difference between the mean sequence and each sample was estimated, and the rate was estimated for each sample. The distribution was slightly distorted (Extended Data Fig. 5), and hence, the representative value was found as the median.

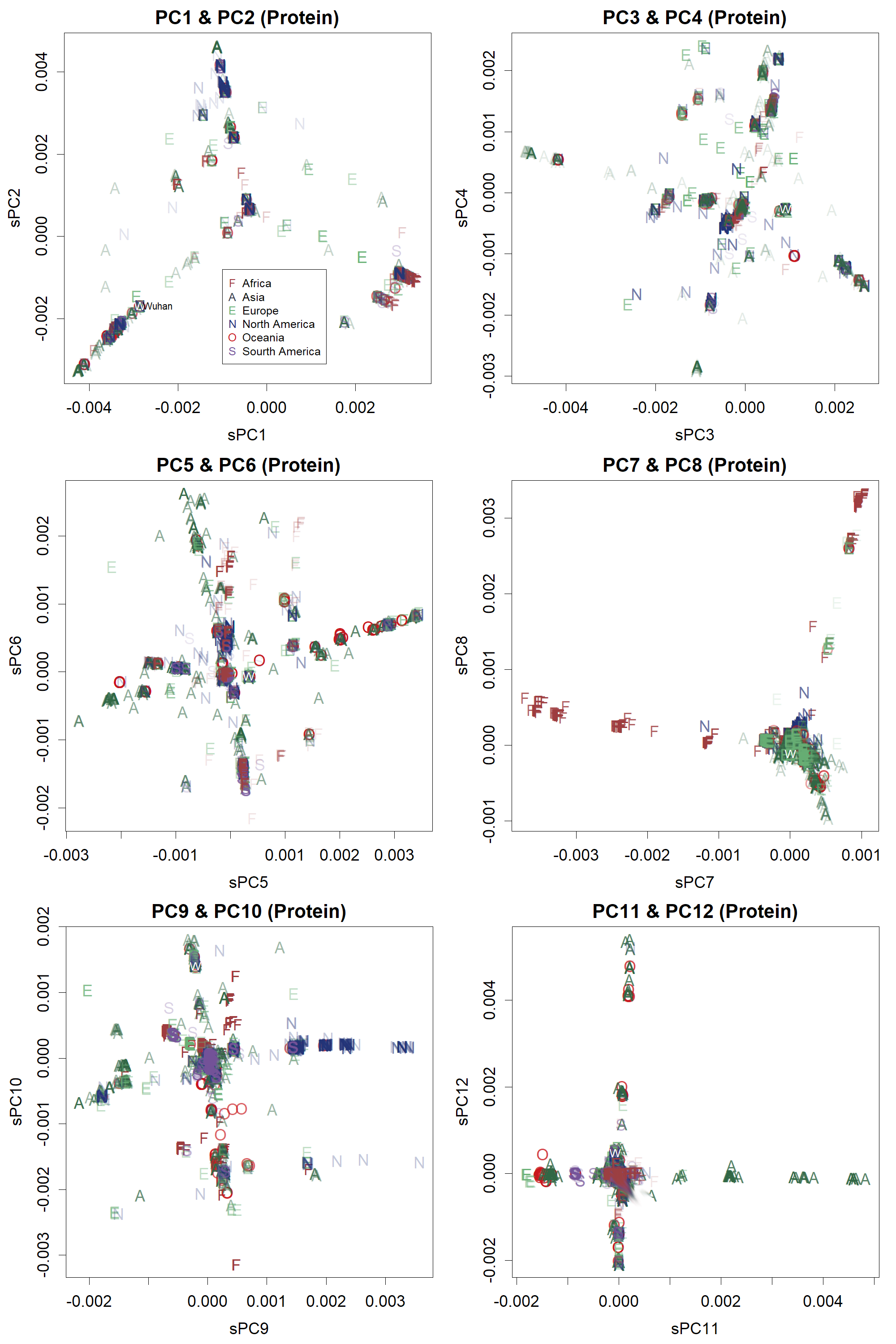

**Extended Data Fig 1:** Principal components of SARS-CoV-2. A thousand samples are selected and shown from each continent. Data are estimated from amino acid sequences.

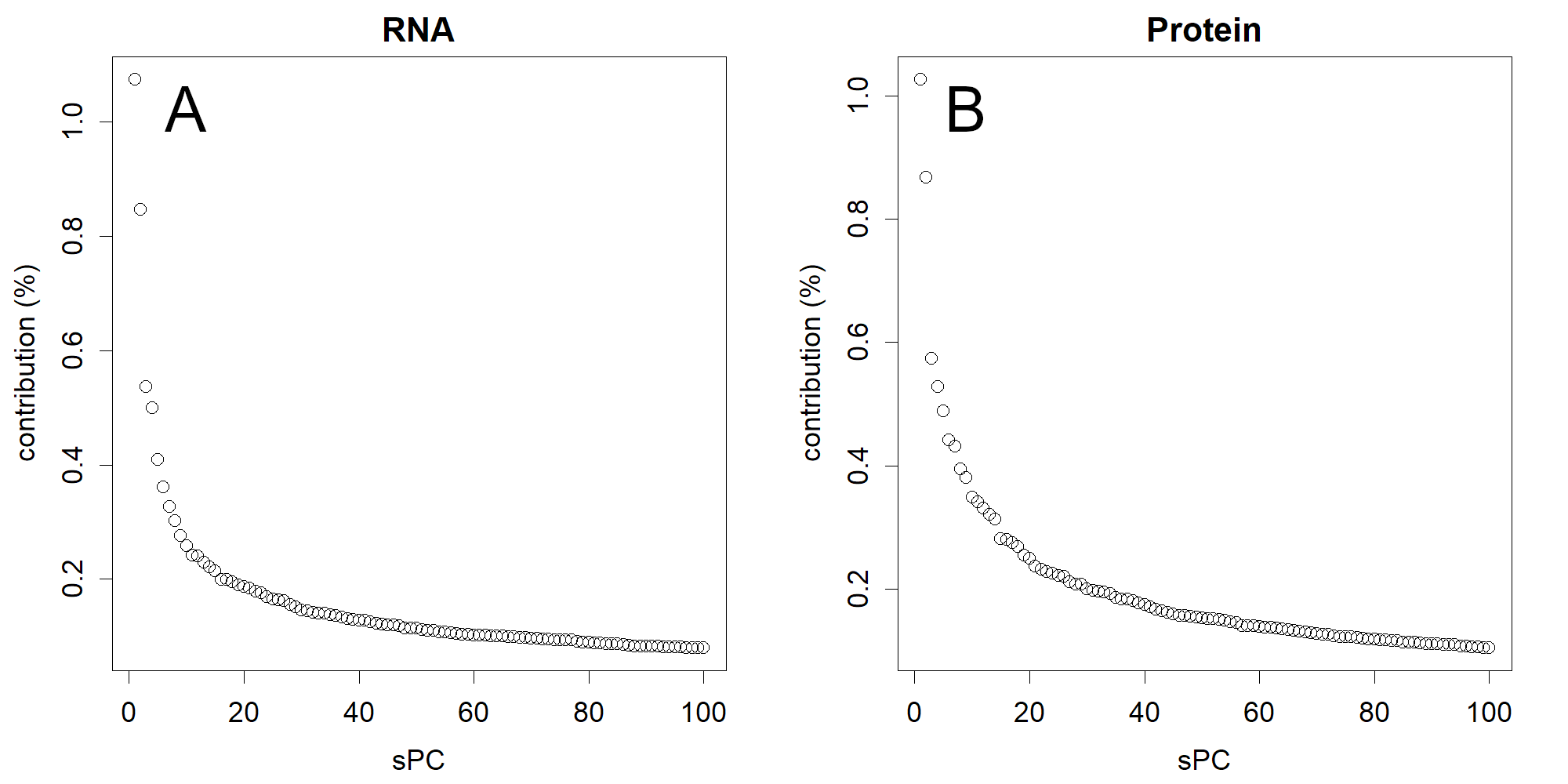

**Extended Data Fig. 2:** Principal component analysis (PCA) contributions. **A.** RNA, **B.** Protein. The superiorities of PC1 and 2, which separate many samples, are clear. In fact, these are very high scores for 6,000 dimensions.

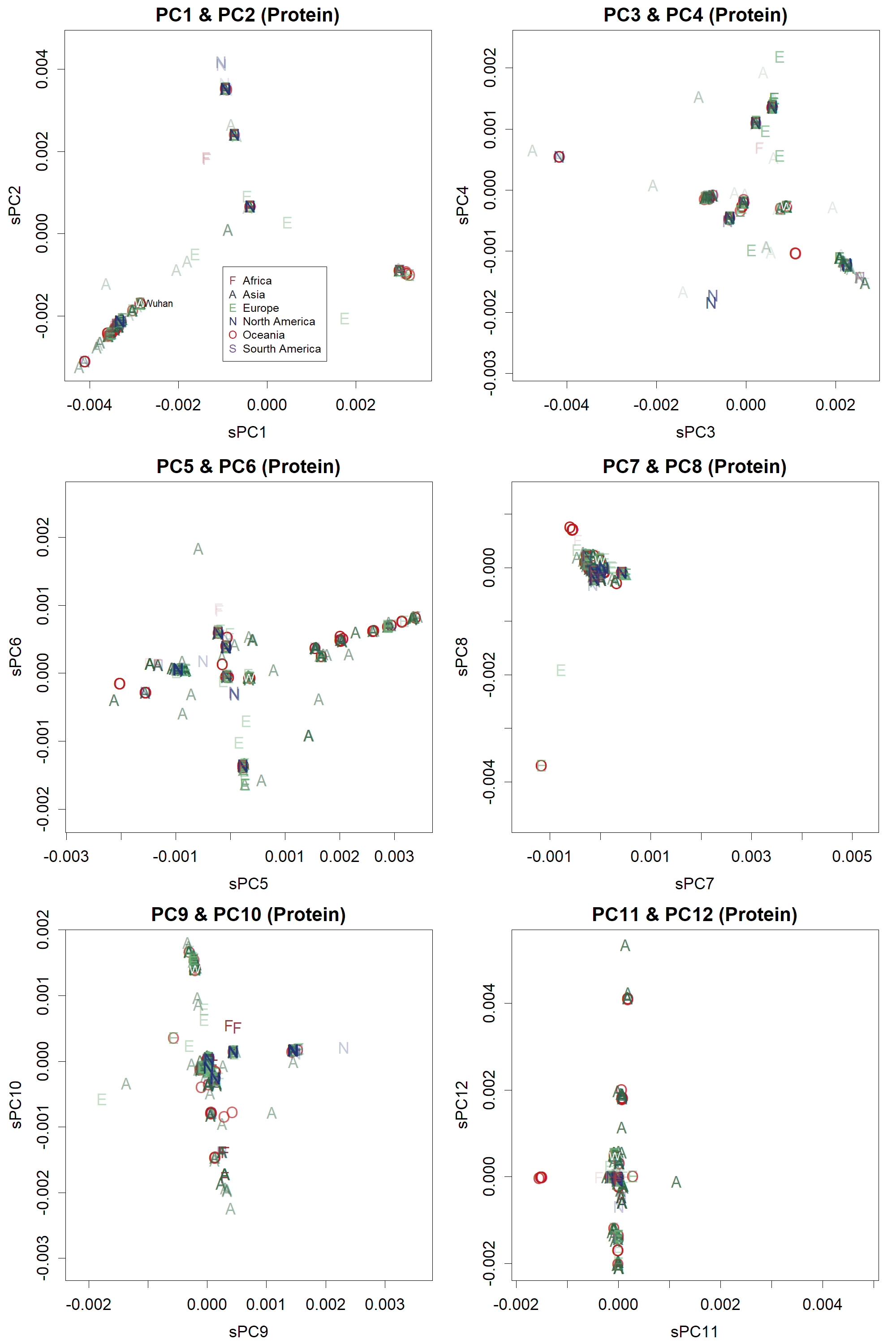

**Extended Data Fig. 3:** Principal component analysis (PCA) of amino acid sequences. Samples collected until the end of March are presented. It is clear that PC1 and PC2 are complete, but the lower axes lack samples that are characteristic of each axis; those have not appeared yet. See Fig. 1 for the full set of amino acid sequences collected until September.

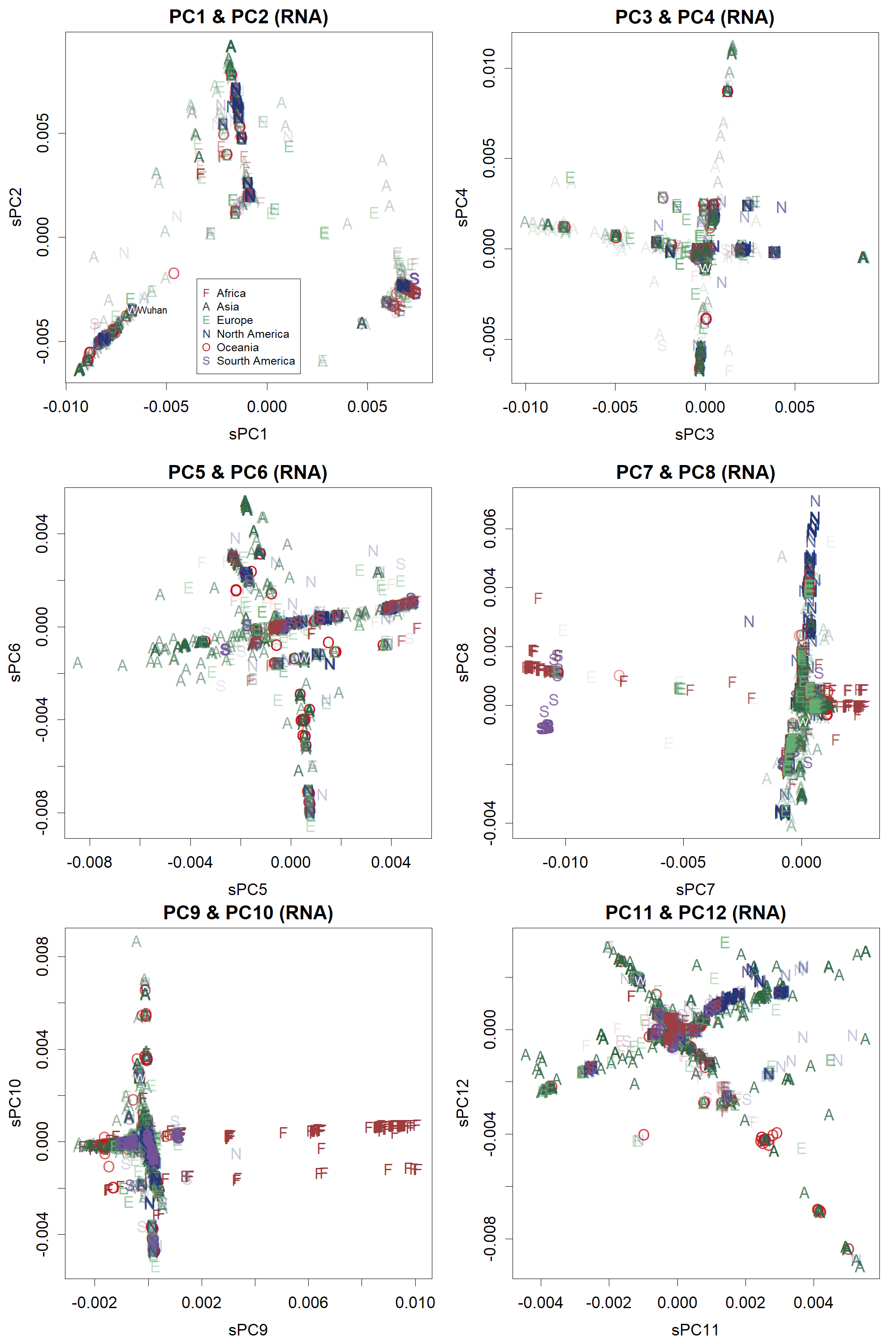

**Extended Data Fig. 4:** Principal components of SARS-CoV-2. A thousand samples are selected and shown from each continent. Data are estimated from nucleotide sequences.

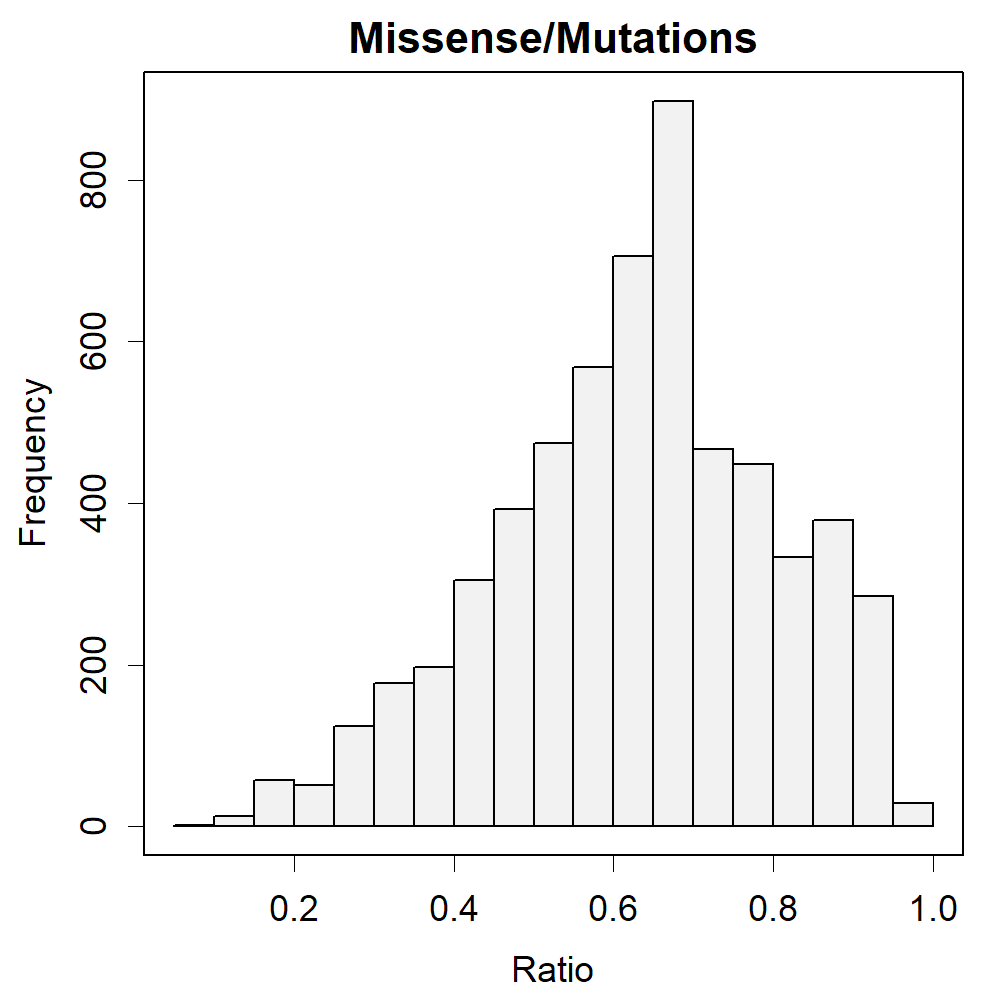

**Extended Data Fig. 5:** The ratio of missense mutations found in SARS-CoV-2 compared with the average sequence. Ratios found in comparison between each sample and the average sequence data are the rate of differences that affect the amino acid sequence.

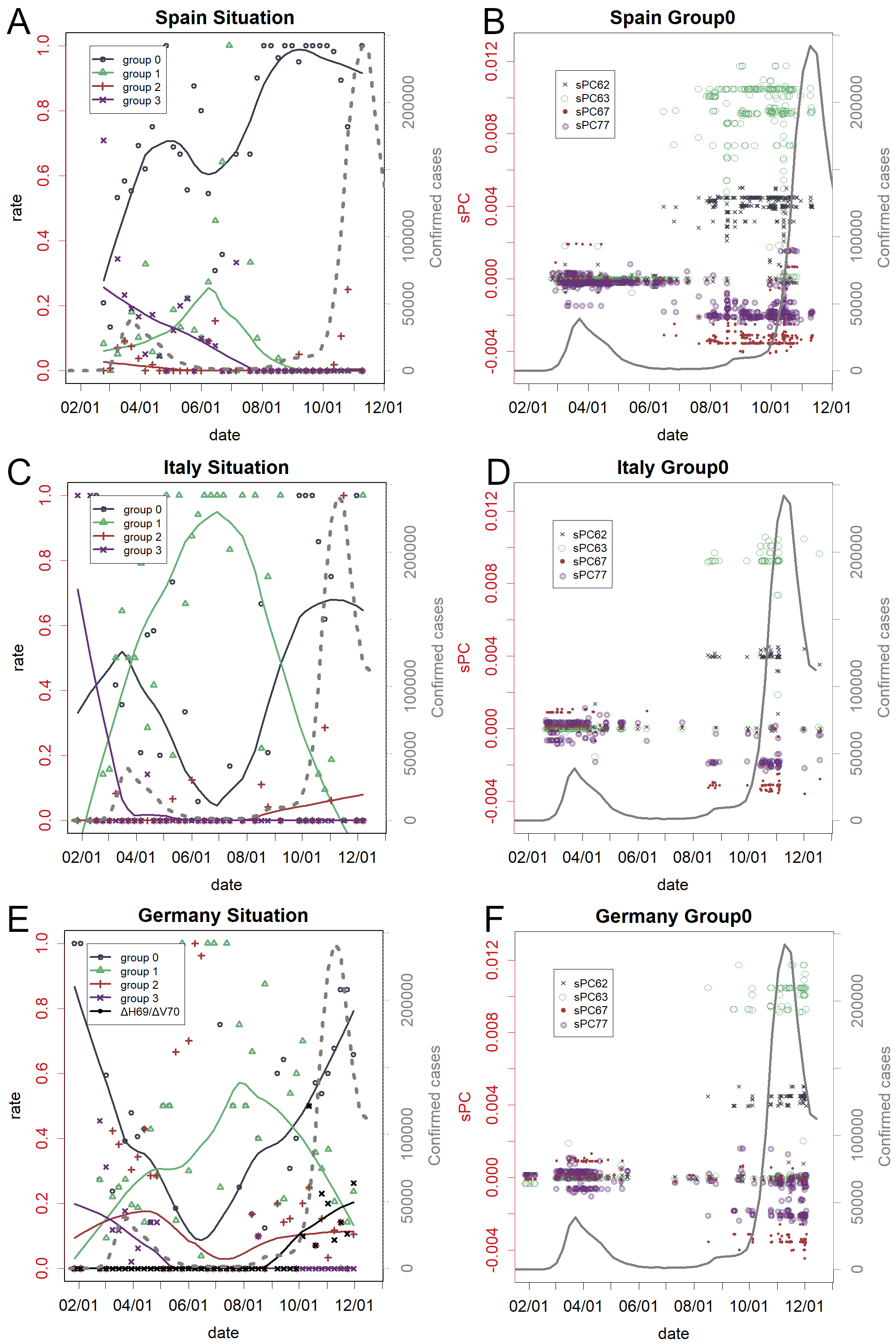

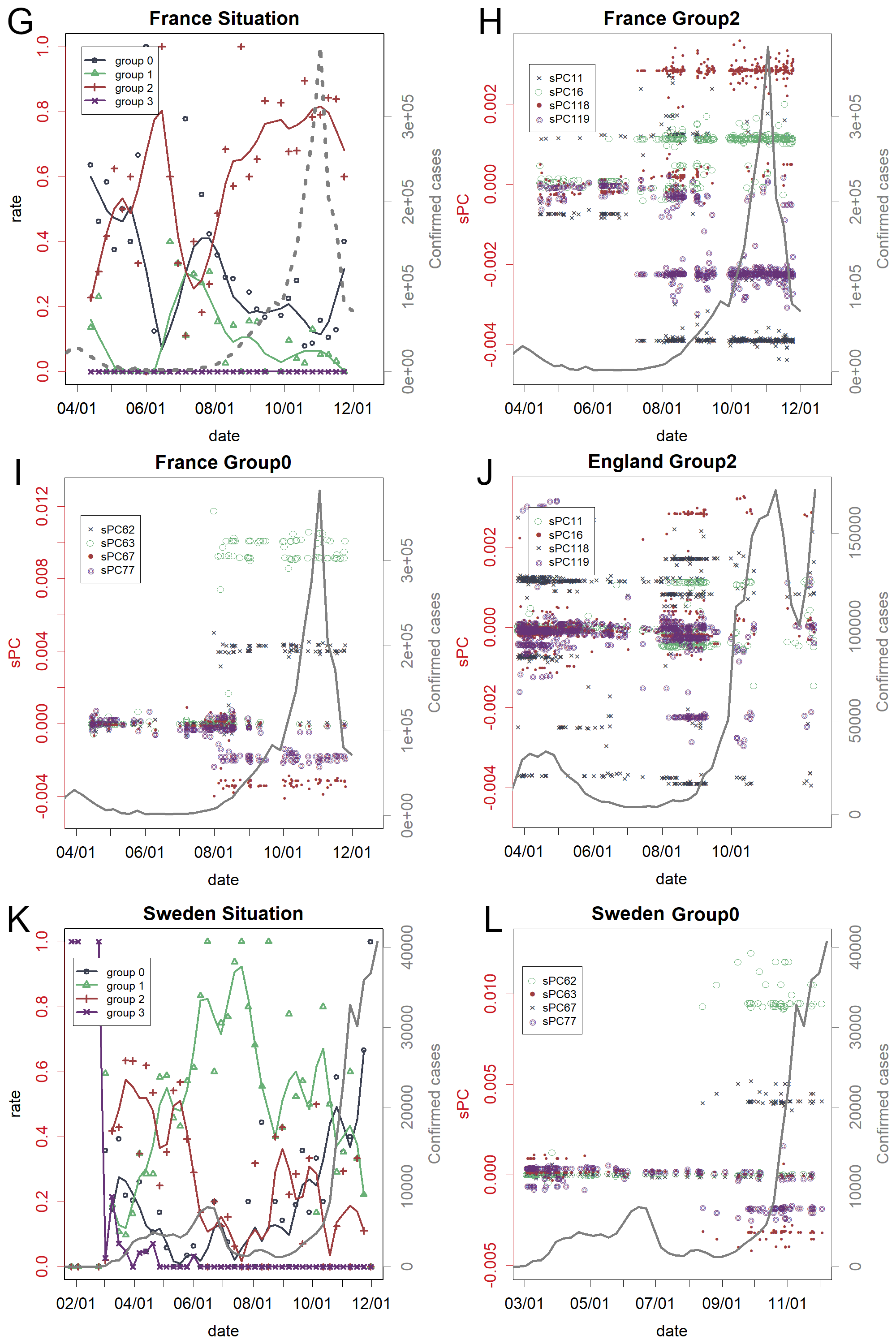

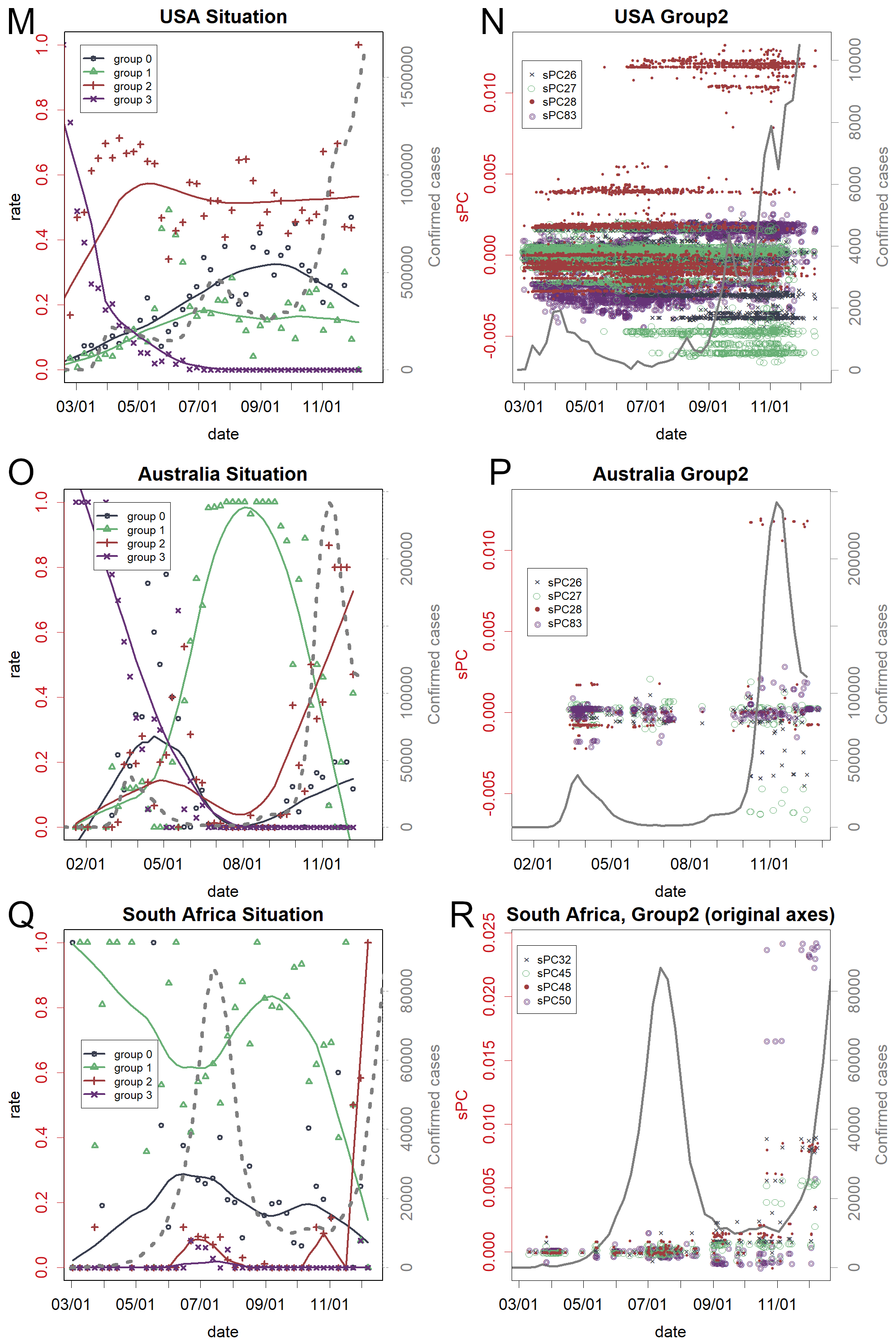

**Extended Data Fig. 6:** Extension for Fig. 2. **A** and **B**: Spain; **C** and **D**: Italy; **E** and **F**: Germany; **G**-**I**: France; J: group 2 of England; **K** and **L**: Sweden; **M** and **N**: USA; **O** and **P**: Australia; **Q** and **R**: South Africa. Panels B, D, F, I, and L presented the pattern of the pan-European variant. The pattern appeared in H and N also appeared in J and P, respectively.

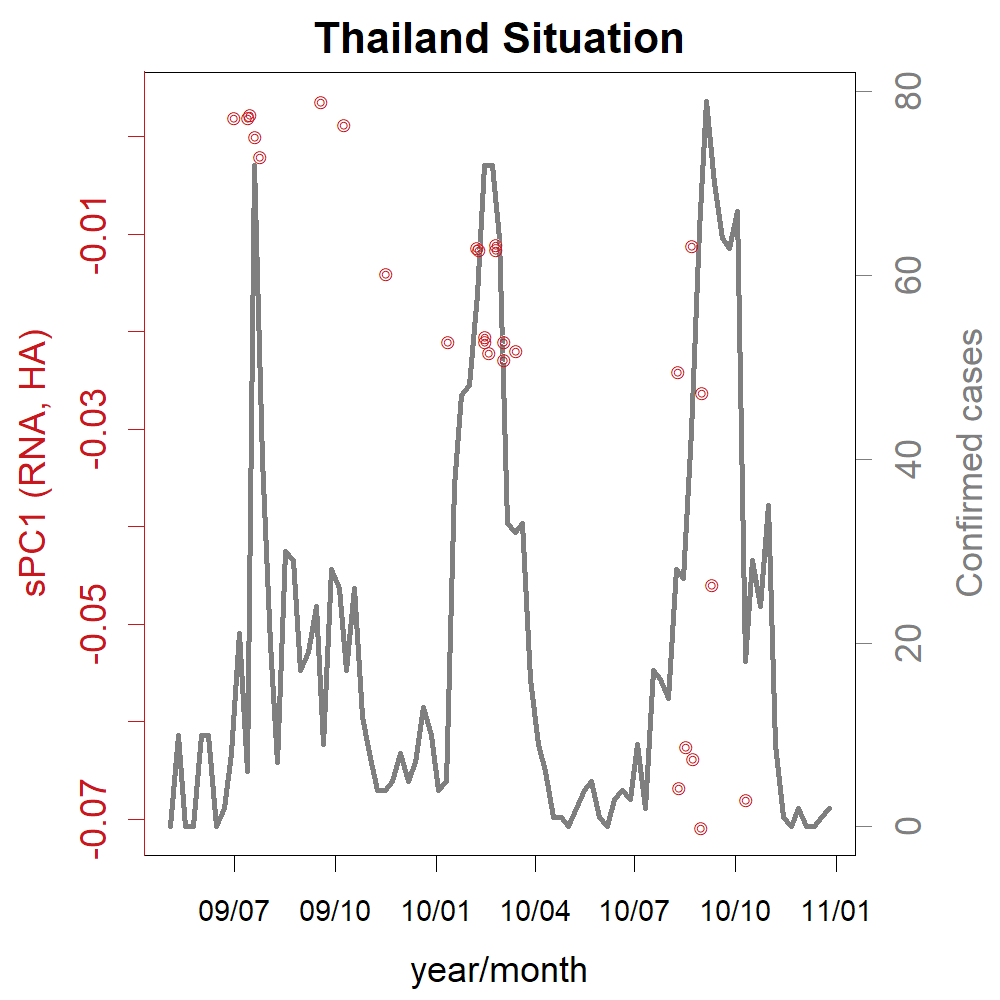

**Extended Data Fig. 7:** Pdm09 influenza in Thailand. The first peak of cases is much smaller than that in other countries. In the second peak, there are no amino acid changes; however, the rate of missense mutation increases in the third peak. The axes have been identified only within this season of the country; otherwise, these small differences will be neglected among the larger variations in the influenza virus.

| SARS-CoV-2 | Example of Mutations |
| --- | --- |
| Austraria (group 1), 4/1 - 8/20 | ORF1ab 300, I/F;S 478, S/N; 1069, V/F |
| France (group 2), 6/5 - 9/20 | ORF1ab 2702, H/Q; 3087, M/I; 4582, A/S; 5173,  V/L; 5547, K/R; 5590, E/D;  S 478, S/N; N 234, M/I; N 376, A/T |
| pan-Europe (group 0), 3/30-8/15 | ORF1ab 7325 A/V; N 221, A/V; ORF10 31 V/L |
| South Africa (group 1), 5/30-8/3 | ORF1ab 2166, T/I; ORF3 125, R/S |
| USA (group 2), 3/30-8/15 | ORF1ab 3352, L/D; 6060, N/D; 7020, R/C; S 173, G/V;  ORF8 25, S/L; N 67, P/S; 200, P/L |
| England (group1), 9/1-11/1 | ORF1ab 2502, I/T; 4242, M/I; 5118, V/I; 5620, H/Y; 5928, A/S; S 69, I/-; 70, H/-; 71, V/-; 440, N/K |
| South Africa (group 2), 9/1-11/1 | ORF1ab 1656, K/N;3354, K/R;  S 81, D/A;243, L/-;244, A/-;245, L/-;246, H/-;418, K/N;485, E/K;502, N/Y;701, A/V;  E 72, P/L; N 206, T/I |

**Extended Data Table 1:** Example of mutations. These are frequent mutations between the peaks. ORF1ab 2702, H/Q: the 2702^th^ residue of ORF1ab protein that changed from H to Q.
